# Near-Zero Missed Cleavages with a High-Fidelity Recombinant Arg-C Zero for Mass Spectrometry-Based Proteomics

**DOI:** 10.64898/2026.05.28.728370

**Authors:** Cristina Hernandez-Rollan, Jonas Damgaard Elsborg, Elisa Le Boiteux, Yang Lu, Karishma Patel, Ivan Ahel, Ole N. Jensen, Tanveer Singh Batth, Jesper Velgaard Olsen

## Abstract

Proteolytic digestion remains a critical step in bottom-up proteomics workflows, with enzyme specificity and efficiency directly impacting peptide identification and protein sequence coverage. Here, we present the comprehensive characterization of Arg-C Zero, a recombinant arginyl endopeptidase derived from *Porphyromonas gingivalis* that exhibits exceptional fidelity in cleaving specifically at the C-terminus of arginine residues. Unlike conventional serine proteases such as Trypsin, Arg-C Zero utilizes a histidine-cysteine catalytic dyad mechanism, achieving near-zero missed cleavage rates (>99% efficiency) under standard proteomics conditions. Through systematic evaluation using HeLa protein extracts, we demonstrate that Arg-C Zero maintains consistent performance across varying digestion times. The enzyme shows robust activity across a broad pH range and tolerates up to 4M urea, making it ideally suitable for a diverse range of proteomics sample preparation workflows. While Trypsin/LysC combinations remain superior for comprehensive proteome coverage, Arg-C Zero offers unique advantages for applications requiring high specificity and reproducible arginine-specific cleavage patterns, particularly for analysis of post-translational modifications (PTMs). Here, we demonstrate how Arg-C Zero aids comprehensive mapping of histone PTMs, and when used in low-pH workflows help preserve labile ADP-ribosylation sites, expanding the analytical capabilities of mass spectrometry for characterizing these challenging modifications. The enzyme’s resistance to proline-adjacent cleavage sites and compatibility with standard mass spectrometry buffers position it as a valuable addition to the proteomics enzyme toolkit.

## Introduction

Proteolytic digestion represents a fundamental step in bottom-up proteomics workflows, where the selection of appropriate proteases directly influences proteome coverage, peptide identification rates, and the overall success of mass spectrometry (MS)-based protein analysis. Traditional workflows predominantly rely on Trypsin, which cleaves at the C-terminus of lysine and arginine residues (*1*), providing predictable peptide lengths and charge states favorable for electrospray ionization and peptide sequencing by MS/MS. However, the reliance on single-enzyme digestion can limit sequence coverage, particularly for proteins with few tryptic cleavage sites or those containing post-translational modifications (PTMs) that inhibit proteolysis.

The introduction of alternative proteases such as Lys-C, Asp-N, and Glu-C have expanded the toolkit available for comprehensive proteome characterization. These enzymes offer complementary cleavage specificities that can enhance overall protein sequence coverage when used in combination or as standalone digestion strategies(*2*). Among these alternatives, arginyl endopeptidases (Arg-C) offer complementary advantages by exclusively targeting arginine residues, generating longer peptides ideal for isoform discrimination and PTM analysis (*3*). In proteomics, the primary organisms and sources associated with Arg-C proteases have been *Clostridium histolyticum* (clostripain) (*4*) (*5*) *Lysobacter enzymogenes* (Endoproteinase Arg-C) (*6*) and mouse submaxillary gland (tissue kallikrein) (*7*). Clostripain is a cysteine-based arginyl endopeptidase that, despite its specific activity towards Arginine, displays inconsistent activity and higher rates of miss cleavages (*8*, *9*). *L. enzymogenes* and tissue kallikrein are serine-based arginyl endopeptidases that, while more stable under standard conditions, exhibit lower specific activity toward arginine residues and broader substrate tolerance, including partial cleavage at lysine residues, which compromises cleavage fidelity in complex proteomes (*10*). A newer type of Arg-C enzyme comes from the gram-negative anaerobic bacterium associated with periodontal disease *Porphyromonas gingivalis. P. gingivalis*, produces several virulence factors, including arginyl endopeptidases known as gingipains (*11*), which have been detected as highly expressed in metaproteomics studies of periodontal health status (*12*). *P. gingivalis* Arg-C displays markedly higher specific activity and selectivity towards the C-terminus of arginine residues compared to both clostripain and the serine-based Arg-C enzymes, cleaving with high efficiency and retaining activity under denaturing conditions compatible with standard proteomics workflows (*13*). The Arg-gingipain (RgpB) from *P. gingivalis* has been extensively characterized for its role in bacterial virulence (*14*) (*15*), but its potential as a proteomics tool has only recently been explored.

Promega’s Arg-C Ultra has demonstrated robust activity across a pH range of 5.0-9.0 and tolerance to denaturing conditions, making it suitable for analysis of difficult-to-digest proteins and preservation of labile post-translational modifications (*16*). Similarly, clostripain-derived Arg-C has been shown to efficiently cleave arginine sites followed by proline, a limitation of Trypsin that affects approximately 5% of all potential cleavage sites (*1*) (*9*). These advances underscore the growing importance of arginine-specific proteases for specialized proteomics applications. One notable example is histone PTM analysis, as these proteins are highly enriched in lysine residues. This lysine-rich composition, together with the presence of PTMs at these sites that can block tryptic cleavage, results in peptides of variable length that can complicate LC-MS/MS analysis, and often requires chemical derivatization (*e.g.* Prop/Prop or Prop/PIC) (*17*) (*18*). These challenges highlight the need for alternative proteases capable of generating peptides suitable for histone PTM analysis without extensive chemical modification.

Here, we present the characterization of Arg-C Zero, a recombinant arginyl endopeptidase derived *P. gingivalis* RgpB protein for high-fidelity proteomics applications. We demonstrate its utility through systematic evaluation of digestion efficiency, sequence coverage, and comparative analysis with established proteolytic workflows. Our findings reveal that Arg-C Zero achieves exceptional cleavage fidelity (>99%), maintains activity across a wide range of experimental conditions, and provides complementary peptide populations that enhance coverage of arginine-rich protein regions underrepresented in tryptic digests. Moreover, unlike Trypsin, which exhibits reduced cleavage efficiency at lysine and arginine residues adjacent to proline, the Arg-C Zero protease can maintain activity towards proline-directed arginine sites, providing more complete digestion profiles. Finally, we demonstrate two applications of Arg-C Zero for PTM proteomics for which it is ideally suited: expanding the sequence coverage of lysine-rich regions, like histones, and preservation of alkaline-labile PTMs, such as ADP-ribosylation, using an augmented Arg-C Zero workflow under mild acidic conditions.

## Materials and Methods

### Arg-C Zero protease preparation

Lyophilized Arg-C Zero, a recombinant version of the *P. gingivalis* RgpB gingipain arginyl endopeptidase was provided by KPL ApS (catalog no. KPL0146, Copenhagen, Denmark, (*19*). For all digestion experiments, Arg-C Zero was reconstituted in water immediately prior to use and added at an enzyme-to-substrate ratio of 1:50 (Enzyme: Substrate) unless otherwise specified.

### Protein Substrates

Bovine serum albumin (BSA, Fraction V, ≥96% purity, Sigma-Aldrich #A7906) was used as a model protein to assess cleavage specificity and sequence coverage. Immortalized human epithelial cervix carcinoma adherent cells (HeLa) were prepared from 500 cm^2^ dishes in DMEM (Gibco, Waltham, MA, USA) media containing 2 mM L-glutamine supplemented with 10% fetal bovine serum (Gibco, Waltham, MA, USA), 100 U/mL penicillin (Life Technologies, Carlsbad, CA, USA), and 100 μg/mL streptomycin in a 37 °C incubator supplemented with 5% CO2. Cells were harvested by Trypsinization, washed three times with ice-cold phosphate-buffered saline (PBS), and lysed in boiling lysis buffer (4% SDS, 100 mM Tris-HCl pH 8.5, 5 mM TCEP, 10 mM chloroacetamide) at 95°C for 10 minutes with vigorous shaking (1000 rpm). Lysates were sonicated using a probe sonicator (3 cycles: 30 seconds on, 30 seconds off, 40% amplitude) to ensure complete cell disruption and DNA fragmentation. Protein concentration was determined using the tryptophan fluorescence assay to ensure accurate quantification. Aliquots were stored at -20°C until use.

### Cell culture and histone extraction

Unless otherwise specified, all cell culture reagents were obtained from Gibco (Thermo Fisher Scientific, Waltham, MA, USA). HepG2/C3A hepatocellular carcinoma cells were cultured in DMEM containing 1 g/L glucose supplemented with 10% fetal bovine serum (Sigma-Aldrich, St. Louis, MO USA), 2 mM GlutaMAX, 1X MEM Non-Essential Amino Acids, and penicillin-streptomycin (100 U/mL and 100 µg/mL, respectively) in a 37 °C humidified incubator with 5% CO_2_. Cells were treated with 15 µM GSK-J4 (MedChemExpress, Monmouth Junction, NJ, USA) or an equivalent volume of DMSO for 24 hours, harvested by Trypsinization, washed twice with PBS, and snap-frozen in liquid nitrogen.

Histone extraction was performed using an acid extraction protocol adapted from Garcia *et al.* (2008). Briefly, frozen cell pellets were resuspended in nucleus isolation buffer (15 mM Tris-HCl pH 7.5, 60 mM KCl, 11 mM CaCl_2_, 5 mM NaCl, 5 mM MgCl_2_, 250 mM sucrose, 1 mM dithiothreitol, 10 mM sodium butyrate, 0.1% Igepal) supplemented with protease and phosphatase inhibitors (cOmplete™ and PhosSTOP from Roche, Basel, Switzerland). Nuclei were isolated by centrifugation (1,000 g, 5 min) and washed twice with nucleus isolation buffer without Igepal. Histones were extracted with 0.2 M H_2_SO_4_ for 1 hour under gentle agitation, and the supernatant was collected after centrifugation (20,000 g, 5 min). Histones were precipitated with trichloroacetic acid (20% final concentration) overnight, recovered by centrifugation (20,000 g, 15 min) and washed once with 0.1% HCl in acetone and twice with pure acetone. Pellets were air-dried and resuspended in ultrapure H_2_O. All steps were performed at 4°C.

Histone purity was assessed by SDS-PAGE and protein concentration was determined using a NanoPhotometer N60 (Implen GmbH, München, Germany).

### Protein purification and *in vitro* ADP-ribosylation assay

Recombinant human PARP10 ART domain (residues 809–1017) was purified from *E. coli*, essentially as previously described (*20*). The protein was diluted to 2 µM in ice-cold NAD^+^ reaction buffer (25 mM HEPES pH 7.5, 150 mM NaCl, 0.5 mM TCEP, 5 mM MgCl_2_), and then incubated with 50mM NAD^+^ (Promega) at 37 °C for 30 min. After the ADP-ribosylation reaction was completed, the sample was flash-frozen and stored at -80 °C until further processing.

### Sample Preparation and Digestion Protocols

Protein samples were prepared using the Protein Aggregation Capture (PAC) method as previously described (*21*). Briefly, 10 μg of protein was mixed with hydroxyl magnetic beads from MagReSyn, (ReSyn Biosciences), or silica-based beads from Chemicell (Chemicell GmbH (Berlin, Germany) and proteins were aggregated by addition of acetonitrile to a final concentration of 70%. Beads were washed twice: once with 100% acetonitrile and once with 70% ethanol. Following washes, beads were resuspended in 50 mM HEPES pH 8.5 digestion buffer containing 10 mM DTT. Arg-C Zezo was added at a 1:50 enzyme-to-substrate ratio (w/w) unless otherwise specified. Digestions were performed in a thermomixer at 37°C with agitation (1000 rpm) for overnight digestion (16 hours) as specified in individual experiments. Trypsin was purchased from (Sigma-Aldrich) and used at a ratio of 1:50 for DDA experiments or 1:100 for DIA experiments.

For phosphoproteomics analysis, 400 ng of digested peptides were desalted using C18 Sep-Pak 96-well plates (40 mg sorbent, Waters) and eluted directly into a KingFisher plate with 75 µL of 80% ACN. Phosphopeptide enrichment was performed on a KingFisher Flex robot using Zr-IMAC HP magnetic beads (ReSyn Bioscience) at a 1:2 peptide-to-beads ratio. The desalted peptides were brought to a final volume of 200 µL with an adjusted loading buffer (80% ACN, 8% TFA, 0.6 M glycolic acid) to normalize for the elution volume from the desalting step. Enrichment was conducted as previously described (*22*, *23*), and the enriched phosphopeptides were transferred to Evotips for LC-MS/MS analysis.

To assess tolerance to denaturing conditions, parallel digestions were performed in the presence of varying concentrations of urea (1M, 2M, and 4M) and guanidinium chloride (0.5M, 1M, and 1,5M). For pH optimization studies, digestions were conducted in buffers adjusted to pH values ranging from 4.5 to 8.5 using 50 mM HEPES. Following digestion, reactions were quenched by addition of trifluoroacetic acid (TFA) to a final concentration of 1%. 500ng of the digested peptides were loaded onto Evotips (Evosep) for LC-MS/MS analysis.

For histone PTM analysis, three technical replicates were prepared for each experimental condition (control and GSK-J4-treated cells). 100ng of histones were digested in 80mM EPPS pH 8 digestion buffer containing 10 mM DTT using Arg-C Zero at a 1:1 protease-to-substrate ratio for 2 hours at 37°C. Resulting histone peptides were loaded onto Evotips (Evosep Biosystems) prior to LC-MS/MS analysis.

For PARP10 ADP-ribosylation analysis, the sample was diluted 100-fold in low-pH digestion buffer (50mM ACES, pH 6.5), and then digested using Arg-C Zero in a 1:20 protease-to-substrate ratio (w/w) for 1 h at 30 °C. The digest was then diluted with guanidine to a final concentration of 2M in 50mM ACES pH 6.5 to inactivate Arg-C Zero, and then subsequently digested with LysC (Wako) in a 1:20 protease-to-substrate ratio (w/w) for 1 h at room temperature, and finally 1 h at 30 °C. The resulting peptides were acidified to final concentration of 0.5% TFA (v/v) and then purified using low-pH C18 stagetip (*24*). Briefly, the C18 material was activated with methanol, and then equilibrated first with 80% ACN in 0.1% formic acid, and then second in 0.1% formic acid. The sample was loaded by centrifugation, and then washed in 0.1% formic acid and finally eluted with 30% ACN in 0.1% formic acid. The elution was dried to completion using a speedvac (45 °C for 3 h), and then redissolved in 0.1% formic acid.

### LC-MS/MS Analysis

#### Enzyme benchmark

Digested peptides (500 ng per injection) were analyzed by liquid chromatography-tandem mass spectrometry using an Evosep One LC system (Evosep Biosystems) coupled to an Orbitrap Exploris 480 mass spectrometer (Thermo Fisher Scientific) or an Astral analyzer. Peptides were separated on an EV-1109 analytical column (PepSep, 8 cm × 150 μm, 1.5 μm particle size) with an EV-1087 emitter (20 μm inner diameter fused silica) using the 60 samples per day (SPD) gradient method.

The Orbitrap Exploris 480 mass spectrometer was operated in data-dependent acquisition (DDA) mode with the following parameters: spray voltage 1.8 kV, ion funnel RF level 40, and heated capillary temperature 275°C. The Orbitrap Astral mass spectrometer was operated in narrow window data-independent acquisition (nDIA) mode (*25*), in positive ionization mode, using a source voltage of 2000 V and the ion transfer tube temperature at 275°C.

#### Histone PTM analysis

Histone peptides (100ng) were analyzed by liquid chromatography-tandem mass spectrometry using an Evosep One LC system (Evosep Biosystems) coupled to a timsTOF Ultra2 mass spectrometer (Bruker Daltonics). Peptides were separated on an Aurora Rapid analytical column (IonOpticks, 5cm × 75µm, 1.7µm particle size) with integrated emitter (compatible with CaptiveSpray Ultra sources) using WhisperZoom80SPD Evosep method. The column was maintained at 50°C using Bruker’s Column Toaster.

The timsTOF Ultra2 mass spectrometer was operated in data-dependent acquisition parallel accumulation–serial fragmentation (DDA-PASEF) mode with a spray voltage of 1.6 kV and a source temperature of 200°C. All spectra were acquired with a m/z range of 100-1700, and ion mobility separation was performed over a range of 0.65–1.25 V.s/cm² (1/K_0_) with an accumulation time of 100 ms and a ramp time of 100 ms. Each acquisition cycle consisted of five PASEF MS/MS scans, resulting in a total cycle time of 0.64 s. Collision energy was linearly ramped as a function of ion mobility from 20 eV at 0.60 V.s/cm² to 59 eV at 1.60 V.s/cm².

#### ADP-ribosylation analysis

Peptides were measured using an Orbitrap Fusion Lumos Tribrid mass spectrometer (Thermo), operated in positive ion mode with a nano-electrospray ion source. The spray voltage was set to 2.0 kV, and the ion transfer tube temperature was 275 °C. Internal calibration was performed using EASY-IC. Peptides were eluted using 20-cm long analytical columns with an internal diameter of 50 μm, packed in-house using ReproSil-Pur 120 C18-AQ 1.9 μm beads (Dr. Maisch). On-line reversed-phase liquid chromatography to separate peptides was performed using an Vanquish NEO system (Thermo), and the analytical column was heated to 40°C using a column oven. Peptides were eluted from the column using a gradient of Buffer A (0.1% formic acid) and Buffer B (80% ACN in 0.1% formic acid), with a total gradient length of 40 min.

Full MS scans were acquired in the Orbitrap over an m/z range of 400–1200 at a resolution of 120,000. Data-dependent acquisition (DDA) was performed with selection of the most intense precursors (intensity threshold 1.1E5) and dynamic exclusion enabled (25 s) to prevent repeated sequencing. Precursors were isolated in the quadrupole with a 2 m/z isolation window.

MS/MS spectra were acquired in the Orbitrap at a resolution of 60,000, with an AGC target of 2E5 (normalized 400%) and a maximum injection time of 180 ms. Fragmentation was performed using ETD with supplemental higher-energy collisional dissociation (EThcD) and calibrated charge-dependent parameters. Only precursor ions with charge states of 3–5 were selected, with up to four MS/MS events per cycle (topN=4).

### Data Analysis

Raw MS data files acquired in DDA mode were processed using MaxQuant software suite version 2.4.13.0 (*26*). Database searches were performed against the UniProt human reference proteome (UniProt reference proteome 2023 release, 20,598 entries). Search parameters included: enzyme specificity set to ArgC/P (cleavage C-terminal to R), up to 3 missed cleavages allowed, carbamidomethylation of cysteine as a fixed modification, and oxidation of methionine and N-terminal acetylation as variable modifications. For semi-specific searches evaluating cleavage specificity, enzyme specificity was set to semi-specific ArgC. Missed cleavage analysis was performed by extracting peptides with 0, 1, or ≥2 missed cleavage sites from the peptide.txt output file. Sequence motif analysis for missed cleavages was conducted using IceLogo software (*27*) to generate positional enrichment logos. Statistical analyses and visualization were performed using R (version 4.3.0) with packages dplyr, ggplot2, tidyr, and biomaRt.

Raw MS data files acquired in nDIA mode were processed using Spectronaut (V.20.3) using the human protein reference database supplemented with the contaminant database from Spectronaut. Cysteine carbamidomethylation was set as a fixed modification, and for variable modifications: N-terminal acetylation and methionine oxidation. Method evaluation was selected, and enzyme specificity was set to ArgC/P with up to 3 miss cleavages allowed.

Bruker .d files were processed using FragPipe version 23.1 and searched against a human histone database downloaded from UniProt, supplemented with the FragPipe (*28*) contaminant database and decoy generation. Digestion enzyme was set to ArgC with up to 1 missed cleavage allowed. Precursor and fragment tolerances were set to 10 ppm and 0.02 Da, respectively. Dynamic modifications included acetylation of protein N-terminus, acetylation/methylation/dimethylation/trimethylation of lysines and phosphorylation of serines and threonines. Peak identities were manually validated based on retention times and inspection of MS/MS spectra. The area under the curve (AUC) was calculated by the software for each histone peptidoform and normalized to the total abundance of peptides with the same primary sequence.

Chromatograms and MS/MS spectra were visualized using Bruker Compass DataAnalysis software version 5.3.

For the ADP-ribosylation specific files, raw MS data were processed using MaxQuant (version 1.5.3.30) with the Andromeda search engine (*29*). Spectra were searched against a custom FASTA database containing the human PARP10 sequence (UniProtID:Q53GL7), supplemented with common contaminants. A reversed decoy database was used to estimate false discovery rates (FDRs).

Enzyme specificity was set to ArgC and LysC as a semi-specific search. The maximum peptide mass was set to 4600 Da, and peptides with a minimum length of 7 amino acids were considered. Carbamidomethylation was not specified as a fixed modification, as alkylation is incompatible with acidic sample preparation. Variable modifications included methionine oxidation, protein N-terminal acetylation, and ADP-ribosylation (+541.06Da mass shift) on C, D, E, H, K, R, S, T, and Y residues, with a maximum of three modifications per peptide. Peptide-spectrum matches and proteins were filtered at a 1% FDR, and site localization was also controlled at 1% FDR. A minimum score of 40 and a delta score of 6 were required for modified peptides (default). After the MaxQuant search, the raw data tables were further filtered at the PSM level using an in-house tool to ensure stringent localisation (loc>90%) of the identified ADPr-sites (https://github.com/Jonaselsborg/Peptide-ADPr-extractor). For visualisation of annotated spectra, we used the IPSA tool (*30*) in combination with MSConvert (*31*) to parse Thermo .raw into mzML format.

## Results

### Biochemical Characterization of Arg-C Zero

The “Rgp2” *Porphyromonas gingivalis* (strain ATCC BAA-308 / W83) gene encodes a protein (Uniprot: P95493) with 5 different domains, however the active form contains only two domains, a 38.8 kDa catalytic domain, and a smaller C-terminus 8.9 kDa IgG-like domain residing beneath the catalytic domain (Figure 1A) as demonstrated previously (RCSB Protein Data Bank ID (PDB): 1CVR, 432 residues) (*32*). The Histidine-Cysteine dyad (H211, C244) catalyzes the amide bond hydrolysis as demonstrated previously using the D-Phe-Phe-Arg-chloromethylketone (FFR-CMK) covalent inhibitor (Figure 1B). The catalytic pocket resides within the catalytic domain at the opposite end of the IgG domain which “cradles” the C-terminus of the catalytic domain. The Aspartic acid (D163) buried within the catalytic S1 pocket, specifically coordinates to the guanidino group of Arginine via Hydrogen bonding (Figure 1B). This catalytic pocket structure very specifically hydrolyses the amide bonds C-terminus to Arginine residues (Figure 1C). We aligned the amino acid sequence of Arg-C Zero to Clostripain, which is another cysteine protease specific for Arginines (*4*, *5*). We found low sequence similarity (∼20%) between the two proteins suggesting high differentiation (Figure 1D). To further assess the structural relatedness between Arg-C Zero (UniProt P95493) and Clostripain (UniProt P09870), we performed pairwise structural alignment using TM-align (Supplementary Figure 1B) (*33*). The same PDB structure identifier for Arg-C Zero (PDB: 1CVR, 432 residues) was used for alignment against the resolved structure of Clostripain (PDB: 9CIP, 459 residues). The resulting TM-score of 0.308 (normalized by the length of Arg-C from *P. gingivalis*, falls at the border of structural randomness, as TM-scores below 0.3 are of no significant structural similarity. These findings are consistent with the distinct evolutionary origins of the two enzymes. Structural superposition of both models was visualized using UCSF ChimeraX (version 1.9), confirming the poor structural overlap and distinct three-dimensional architectures of the two proteases (Supplementary Figure 1A), where Arg-C Zero is displayed in blue colour and Arg-C from Clostripain is in ocher.

**Figure 1.**
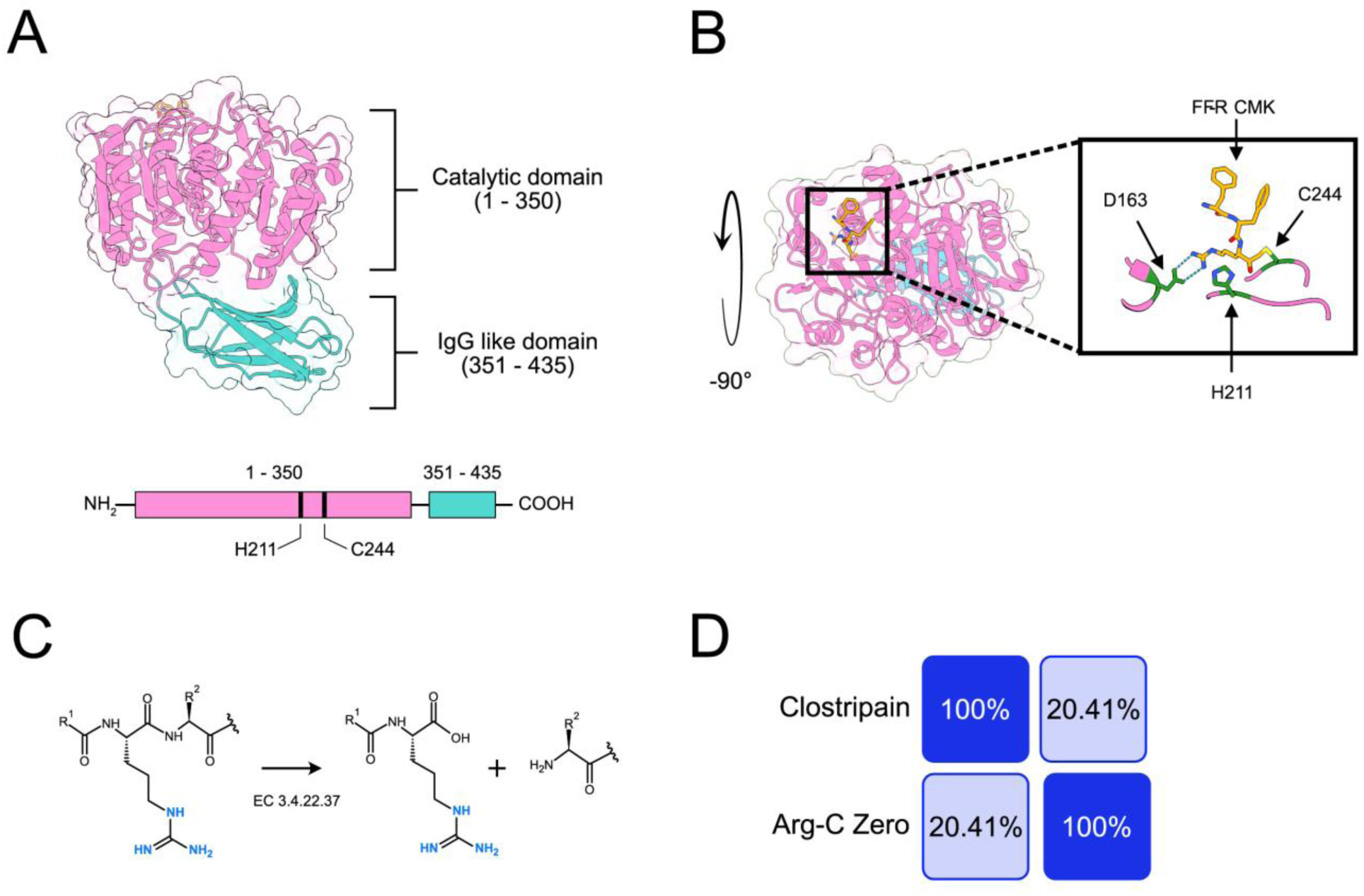
Structural comparison of Arg-C Zero. **(A)** Cartoon representation of the Arg-C Zero three-dimensional structure. The catalytic domain (pink, residues 1-350) and the Ig-G-like domain (teal, residues 351-435) are indicated, highlighting the catalytic residues H211 and C244. **(B)** Active site architecture of Arg-C Zero shown in a -90° rotation view with the inhibitor FFR-CMK bound to the active site. **(C)** Reaction mechanism for Arg-C Zero (EC 3.4.22.37), showing the hydrolysis of the C-terminal side of arginine residues. **(D)** Sequence identity matrix comparing Arg-C Zero and Arg-C from Clostripain, showing that even though both are cysteine enzymes with arginine cleavage specificity, they are structurally distinct, sharing only 20.41% of sequence identity.

#### Effect of Enzyme-to-Substrate ratio on digestion performance

To determine the optimal enzyme concentration for Arg-C Zero, we evaluated the digestion performance across a range of enzyme-to-substrate ratios ranging from 1:25 to 1:500. As expected, performance scaled with enzyme concentration, with the 1:25 ratio yielding the highest identifications: 59,930 peptides and 7,693 protein groups. Performance declined progressively at higher dilutions, and at the 1:500 dilution ratio, Arg-C retained substantial activity, identifying 27,758 peptides, and 5,920 protein groups, approximately 77% of the protein groups identified at 1:25 ratio. In a similar fashion, the missed cleavage showed an inverse relationship with enzyme concentration, in which the optimum of <1% missed cleavage rates were observed for the 1:25 and 1:100 ratio, and increased to 4,2% for the most diluted concentration of Arg-C Zero at 1:500 ratio. This activity performance, even at this 20-fold dilution range suggests that Arg-C Zero is well suited for workflows where enzyme quantity is limited (Figure 2G-I).

**Figure 2.**
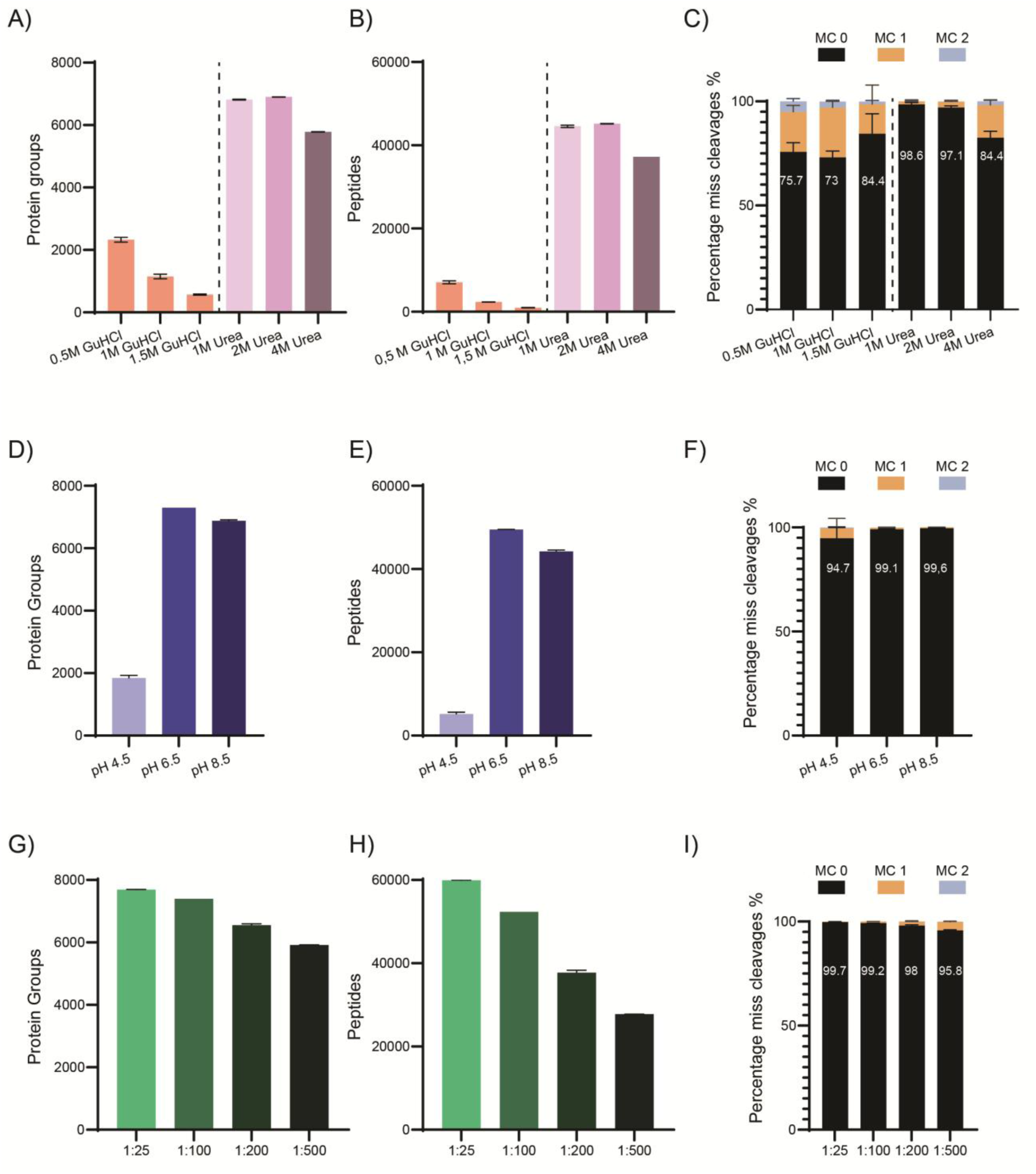
Biochemical characterization of Arg-C Zero digestion conditions in human cell lysate analyzed by LC-MS/MS. Protein groups (A, D, G), unique peptides (B, E, H), and missed cleavage distributions based on peptide intensities (C, F, I) identified across varying digestion conditions. **(A–C)** Effect of denaturant type and concentration. GuHCl (pink, 0.5–1.5 M) and urea (purple, 1–4 M) were compared, with urea at 1–2 M yielding optimal protein group and peptide identifications. The dashed line separates the two denaturant series. MC=0 rates under urea conditions ranged from 84.4–98.6%, with 1 M urea achieving the highest completeness. **(D–F)** Effect of digestion pH. Digestion at pH 4.5 (light blue) resulted in substantially reduced identifications, while pH 6.5 and 8.5 (dark blue) performed comparably, yielding >7,000 protein groups and >44,000 peptides. MC=0 rates improved with increasing pH, reaching 99.1% and 99.6% at pH 6.5 and 8.5, respectively. **(G–I)** Effect of enzyme-to-substrate (E:S) ratio. Ratios from 1:25 (light green) to 1:500 (dark green) were evaluated. Protein group and peptide identifications declined progressively at lower enzyme amounts, with the most pronounced drop at 1:500. MC=0 rates remained high (≥95.8%) across all ratios tested, demonstrating robust cleavage efficiency even at low enzyme concentrations. Error bars represent standard deviation of four technical replicates (n = 4). MC, missed cleavages. Data analyzed by nDIA on Orbitrap Astral analyzer, data analyzed on Spectronaut 20.3.

#### Effect of denaturant agents for digestion with Arg-C Zero

The compatibility of Arg-C Zero with denaturing agents was evaluated using two common chaotropes, guanidinium hydrochloride (GuHCl) and urea, at concentrations relevant to proteomics sample preparation workflows using a protein to enzyme ratio of 1:50 using a HeLa lysate. The two denaturant agents had markedly different effects on Arg-C Zero performance. GuHCl has a high inhibitory effect even at low concentrations, where at 0.5M GuHCl, only 7,070 peptides, and 2,329 protein groups were identified, representing a dramatic reduction compared to standard conditions. Activity was further suppressed at 1.5 M GuHCl, indicating that GuHCl is not compatible with Arg-C Zero digestion and should be avoided or fully removed prior to proteolysis (Figure 2A-C).

In contrast, urea was well tolerated across all tested concentrations up to 4M. Surprisingly, performance peaked at 2M urea (45,171 peptides, 6,902 protein groups), suggesting that mild urea-mediated denaturation conditions may improve substrate accessibility and enhance digestion efficiency. Even at 4M urea, digestion was 79% retained compared to the 2M optimum. These data demonstrate that Arg-C Zero is well suited for urea-based denaturation workflows up to at least 4 M.

#### Effect of pH on Digestion Performance

To find the optimal pH range for Arg-C Zero activity, digestions were performed in 50mM HEPES buffer adjusted to pH 4.5, 6.5, and 8.5. The enzyme exhibited a strong pH dependence, where at pH 4.5 activity was severely impaired, yielding only 5,155 peptides, and 1,840 protein groups. Activity increased dramatically at pH 6.5 with miss cleavages of approximately 2%, and a modest decline was observed at pH 8.5, retaining 90% of the peptide identifications and 94% of the protein groups relative to the 6.5 optimum in activity. However, missed cleavage rates continued to decrease at pH 8.5, reaching approximately 1%, suggesting that while a mild decrease in identifications are generated at alkaline pH, peptides are produced in a more complete cleavage even (Figure 2D-F). The progressive effect of cleaving efficiency at pH 8.5 indicates that the enzyme catalytic mechanism is progressively more efficient at higher pH. Taken together, this data reflects a pH optimum of 6.5 to 8.5 for Arg-C Zero, with 8.5 offering the best balance of activity and efficiency for routine applications.

#### Effect of Reducing Agent on Digestion Performance

To assess the compatibility of Arg-C Zero with common reducing agents used in proteomics sample preparation, we compared dithiothreitol (DTT) and tris(2-carboxyethyl)phosphine (TCEP) (Supplementary Figure S2). Missed cleavage rates were comparably low across all conditions, ranging from 0.3% to 0.9%, indicating that the choice of reducing agent does not substantially compromise Arg-C Zero cleavage efficiency. However, inspection of the total ion chromatograms (TICs) revealed a pronounced late-eluting peak at approximately 20–21 minutes in both TCEP-containing samples, which was entirely absent in DTT-treated samples (Supplementary Figure S2B). Given that DTT provides equivalent digestion efficiency without chromatographic interference, DTT is the recommended reducing agent for Arg-C Zero-based proteomics workflows.

#### Arg-C performance comparison with Trypsin

To benchmark Arg-C Zero against Trypsin under standard bottom-up proteomics conditions, we performed PAC digestions of HeLa cell lysate at an enzyme to protein ratio of 1:50 for data-dependent acquisition (DDA) and 1:100 ratio for data-independent acquisition (DIA) with Arg-C Zero and Trypsin, respectively. We subsequently analyzed the resulting peptide mixtures by both (DDA) and (DIA) mass spectrometry, using three replicates per condition. Raw files were searched against the human Uniprot proteome using MaxQuant (DDA) or Spectronaut (DIA). In DDA mode, Arg-C Zero identified a mean of 9,500 peptides per replicate, compared to 12,400 for Trypsin (Figure 3A). However, despite this difference in peptide count, Arg-C Zero identified 2,467 protein groups compared to 2,333 for Trypsin (Figure 3B). A high overlap of 2076 identified proteins demonstrates that the peptides resulting from Arg-C Zero digestion create a comparable proteome coverage as digestion with Trypsin (Figure 3C). In DIA mode, Arg-C Zero identified substantially lower peptides ∼67,000 per replicate versus Trypsin, ∼154,000 (Figure 3D). However, similar as for DDA, the resulting protein group identifications were comparable between Arg-C and Trypsin with 8,284 and 8,389, respectively (Figure 3E). Venn diagram analysis of the protein identifications from DIA reveals a substantial overlap between the two enzyme digestions in both acquisition modes (Figure 3F). Analyzing the digestion efficiency, Arg-C Zero produces peptides with ∼99% zero miss cleavages (99,4% in DDA, 98,9% in DIA), compared to 76,6% in DDA and 68,5% in DIA for Trypsin (Figure 3G). Analysis of the charge state (z) distribution of detected peptides revealed marked differences between the two enzymes (Figure 3H) While Trypsin generates peptides with doubly charge (z=2, 56,1%), Arg-C Zero generates peptides distributed across higher charge states, with the largest fraction detected at z=3, followed by z=2 and z=4. No singly charged peptides were detected for either enzyme. This shift towards higher charge states for Arg-C Zero is in agreement with the generation of longer peptides bearing multiple ionizable sites. Indeed, after evaluating the amino acid length resulting from the digestion by either Arg-C Zero or Trypsin, we identified that Arg-C Zero produces peptides which are significantly longer (median 15 aa for Arg-C Zero vs 12 aa for Trypsin), reflecting the absence of lysine cleavage sites (Figure 3I). As Arg-C Zero only cleaves after arginine it generates fewer peptides per protein by an in-silico digest compared to Trypsin and hence a smaller database search space. To assess the effect of this, we looked at the run time spent in analyzing the DIA raw files in DIA mode (Spectronaut). While the run time of the raw files analyzed for Trypsin was 69,6 minutes, the raw files for Arg-C Zero were analyzed in 48,48 min, roughly 30% faster (Figure 3J).

**Figure 3.**
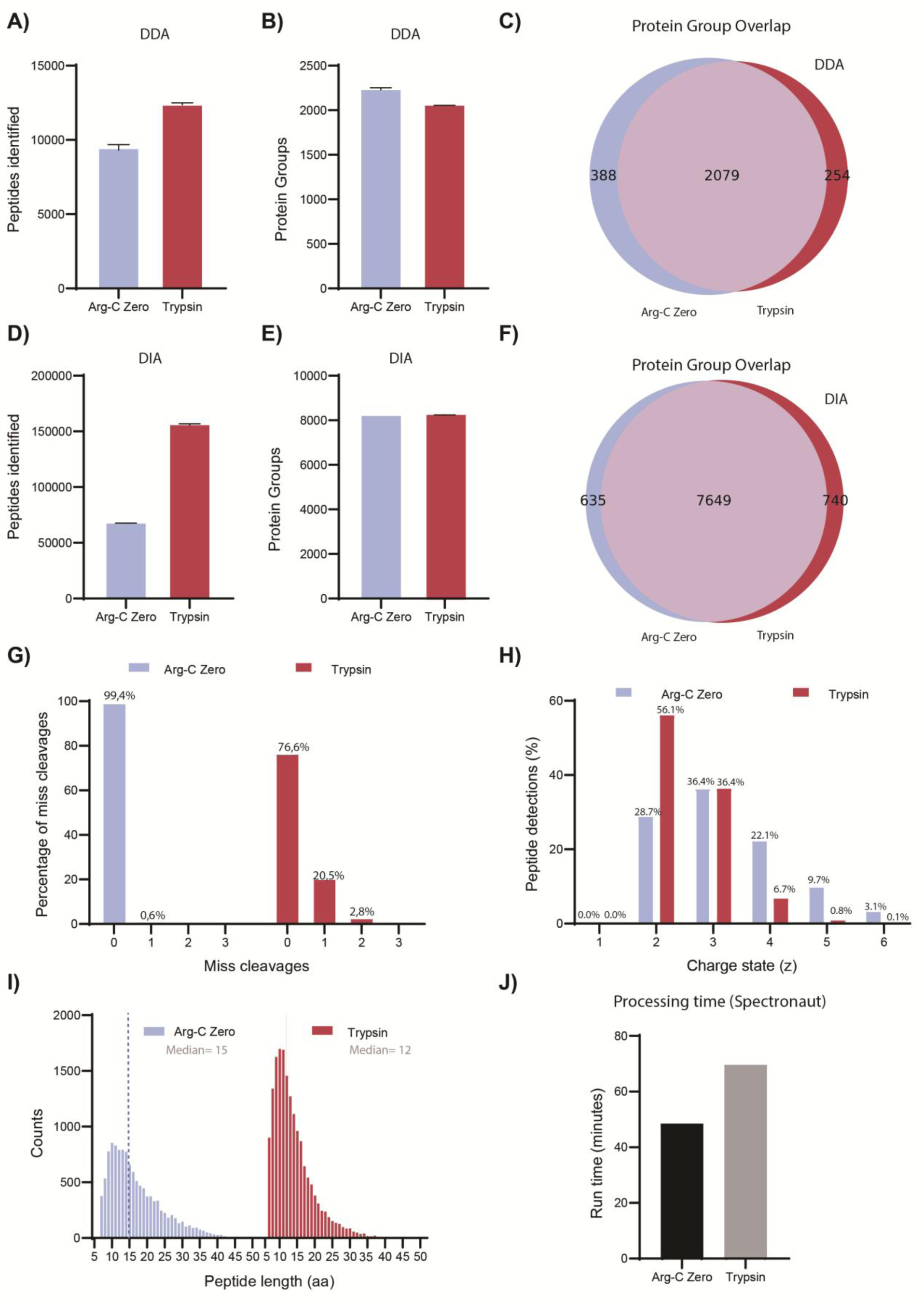
Performance comparison of Arg-C Zero and Trypsin in DDA and DIA mode. **(A)** Mean peptides identified per replicate in DDA mode (n=3). **(B)** Total protein groups identified in DDA mode. **(C)** Venn diagram of protein group overlap between Arg-C Zero and Trypsin in DDA. **(D)** Mean peptides identified per replicate in DIA mode (Spectronaut, n=3). **(E)** Total protein groups identified in DIA mode. **(F)** Venn diagram of protein group overlap between Arg-C Zero and Trypsin in DIA. **(G)** Miss cleavage distribution for peptides identified in DDA. **(H)** Charge state distribution of detected peptides for each enzyme in DDA mode. **(I)** Peptide length distribution for DDA-detected peptides. **(J)** Spectronaut processing time (minutes) for DIA datasets. Enzyme to protein ratio for Arg-C Zero and Trypsin of 1:50 for DDA experiments and 1:100 for DIA.

### Cleavage Specificity and Sequence Context Analysis

To ensure that missed cleavage rates accurately reflect the dominant digestion products rather than the mere presence of low-abundance incomplete cleaved species, the miss cleavage analysis was performed on an intensity-weighted basis, rather than from raw peptide counts (Figure 2C, F, I). For reference, count-based missed cleavage distributions are provided in Supplementary Figure S3).

To comprehensively evaluate cleavage specificity, we analyzed the overnight HeLa digestion data using MaxQuant with semi-specific search parameters to identify all proteolytic cleavage events regardless of specificity. We performed iceLogo analysis (*27*) comparing R-cleaved peptides (positive set, n=14,732) against non-R cleavage sites (negative set= 2,053). The resulting sequence logo revealed a dominant arginine (R) at the P1 position, with a percentage frequency above 100%, demonstrating strict arginine preference (Figure 4A). Consistent with high specificity, analysis of the cleaved peptides revealed a specificity of 98,98% in which all identified peptides were fully cleaved (MC=0, n=16,614), with only 171 peptides (1,02%) having one or more internal uncleaved arginine sites (Figure 4B). Among the miss cleavages events, the majority contained a single miss cleavaged site (MC=1, n=140), with gradually fewer peptides carrying 2, 3, or 4 miss cleavages (MC=2, n=26; MC=3, n=4; MC=4, n=1).

**Figure 4.**
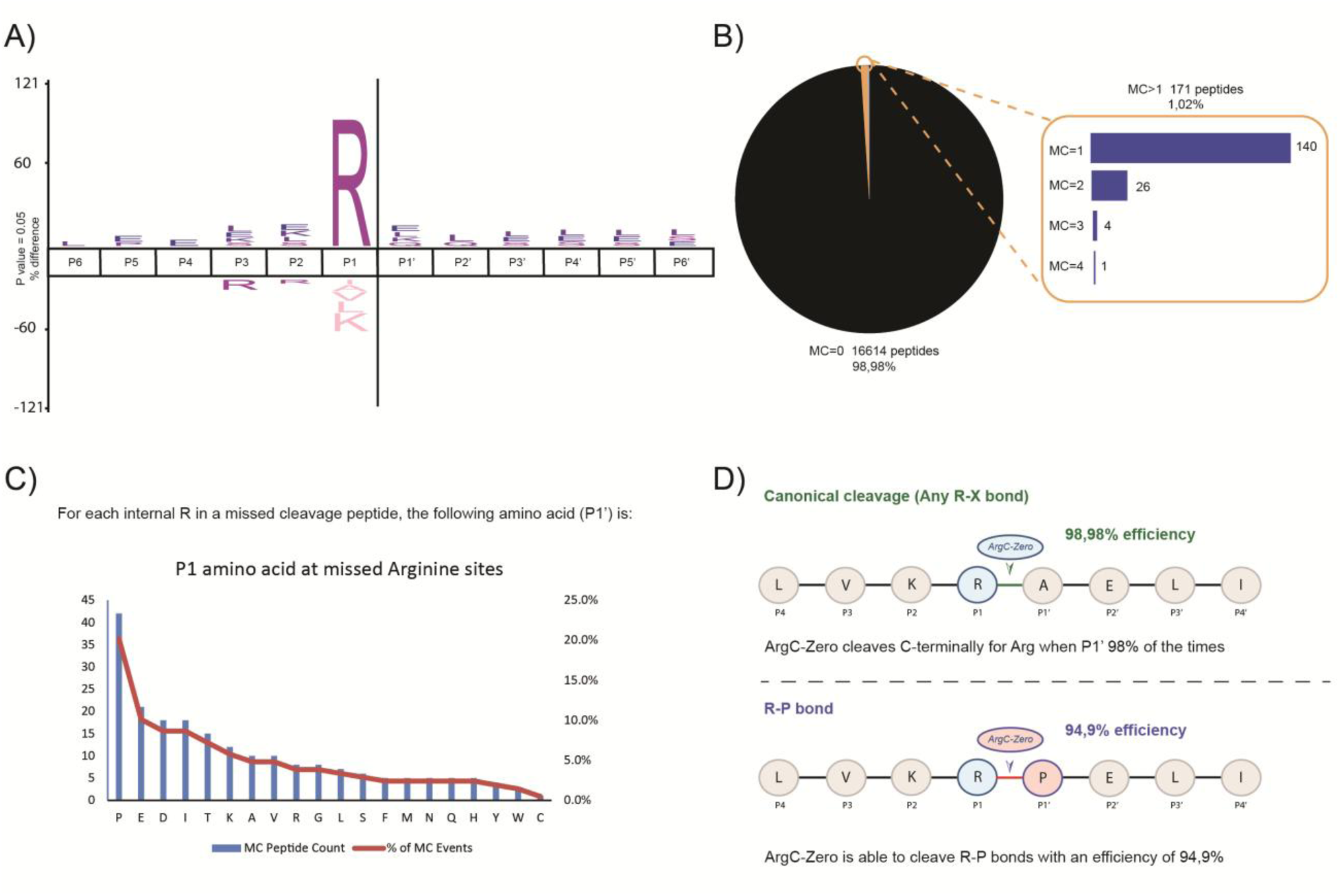
Cleavage specificity and missed cleavage analysis of Arg-C Zero. **(A)** IceLogo analysis of Arg-C Zero cleavage specificity. Amino acid frequencies at P6 to P6’ were evaluated between R-cleaved peptides and non-R cleaved peptides identified in the semispecific database search. Amino acids displayed above the axis are over-represented in the R-cleaved set in comparison to the non R-cleaved set, and amino acids below the axis are depleted. The dominant arginine (R) at P1 confirms strict cleavage specificity. **(B)** Missed cleavage distribution across the data set. **(C)** P1’ amino acid distribution at internal missed arginine cleavage sites. For each peptide carrying at least one miss cleavage, the following residue immediately C-terminal to every internal uncleaved arginine (P1’) was evaluated and their relative miss cleavage rate percentage. **(D)** Schematic illustration of Arg-C Zero cleavage behaviour at R-X and R-P bonds. Upper panel: at canonical R-X bonds (any P1’ residue), Arg-C Zero cleaves with 98,98% overall efficiency.

To identify sequence features that might contribute to missed cleavages of that 1,02%, we looked systematically at the P1’ residue at each internal uncleaved arginine site among all miss-cleaved peptides. For all the amino acids at P1’ position, proline was the most frequent amino acid observed, accounting for approximately 22% of all miss-cleaved events, and 41 events (Figure 4C). Although proline at P1’ is the most frequent driver of miss cleavages, Arg-C Zero is still able to retain 94,9% cleavage efficiency at R-P bonds (Figure 4D), making the global impact of R-P bonds almost negligible, confirming that Arg-C Zero is a robust arginyl endopeptidase across all context sequences.

### Application to Histone Post-Translational Modification Analysis

To evaluate the performance of the enzyme for histone PTM analysis, we analyzed purified histones digested using Arg-C Zero and assessed PTM detection and quantification by LC–MS/MS.

Using Arg-C Zero digestion, we achieved high sequence coverage across all core histones (*e.g*., 82% for histone H3.2 and 71% for histone H4) and detected multiple PTM sites, with 31 and 19 distinct peptidoforms identified on histones H3.2 and H4, respectively (Figure 5A). These PTMs, predominantly localized on the N-terminal tails, include lysine methylation (mono-, di-, and tri-), lysine acetylation, and serine phosphorylation.

**Figure 5.**
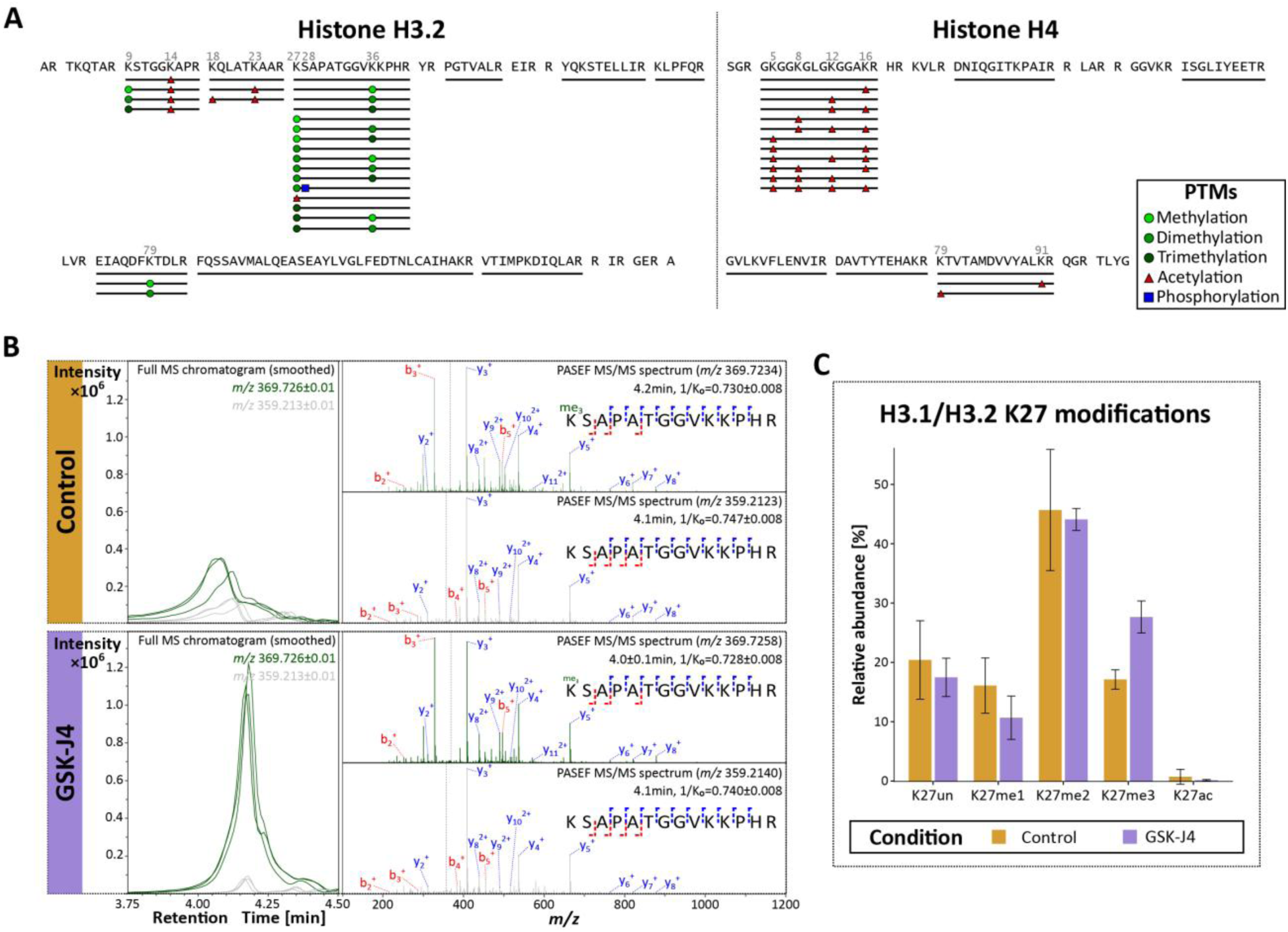
Identification and quantification of histone peptidoforms generated using Arg-C Zero. **(A)** Amino acid sequence coverage of histones H3.2 and H4 following Arg-C Zero digestion. Distinct peptidoforms were mapped onto the protein sequences, and identified PTMs and their position are indicated on the peptides. **(B)** Extracted ion chromatograms (left) and corresponding MS/MS spectra (right) of the H3.1/H3.2 K27-containing peptide in its unmodified (grey) and trimethylated (green) forms in control and GSK-J4-treated cells (n=3 technical replicates). Characteristic b- (red) and y- (blue) fragment ions are shown, demonstrating localization of trimethylation on lysine 27 (+42.047 Da mass shift). **(C)** Relative abundance of H3.1/H3.2 K27 modifications, calculated from the 16 identified peptidoforms, in GSK-J4-treated cells compared to control. Data are presented as mean ± SD. un=unmodified; me1/2/3=mono-, di-, tri-methylated; ac=acetylated.

To further assess whether Arg-C Zero enables accurate detection of biologically relevant PTM changes, we analyzed hepatocellular carcinoma cell culture treated with GSK-J4, a selective inhibitor of the JMJD3 (KDM6B) demethylase that targets trimethylated lysine 27 on histone H3 (H3K27me3), and inspected LC-MS features. Extracted ion chromatograms of the H3.1/H3.2 peptide containing K27 (KSAPATGGVKKPHR) in control and GSK-J4-treated cells demonstrated a marked increase in the trimethylated species (green curves, n=3) upon GSK-J4 treatment, whereas no change was observed for the unmodified species (grey curves, n=3) compared to control cells (Figure 5B, left panel). This observation was consistent across the three technical replicates, demonstrating high reproducibility. MS/MS spectra unambiguously localized the trimethylation to lysine 27, as fragment ions carrying the modification all contained the residue K27, whereas no fragments indicated modification at neighboring lysine K36 (Figure 5B, right panel), confirming that the observed increase in trimethylation in GSK-J4-treated cells occurs specifically at H3K27.

This finding was confirmed by quantitative analysis of the 16 peptidoforms sharing the same amino acid sequence, which revealed that GSK-J4 treatment resulted in a redistribution of K27 peptidoforms characterized by an increase in H3K27me3 levels and a concomitant decrease in H3K27me1, indicating a shift towards higher methylation states at H3K27 (Figure 5C). Together, these results demonstrate that Arg-C Zero enables precise detection and quantification of site-specific histone PTM dynamics.

#### Acid compatible digestion using Arg-C Zero preserves labile PTMs

ADP-ribosylation is a modification catalysed by the PARP-enzyme family (*34*), which involves the transfer of a ADP-ribose (ADPr) onto other targets using NAD^+^ as a substrate. The PARP-family, found in all kingdoms of life, consists of several members with distinct functional roles, including a diverse selection of the amino-acids they modify (*35*, *36*). While ADP-ribosylation of serine is remarkably stable, modification of Glu and Asp form ester-bonds, which are labile with conventional MS sample preparation(*37–39*). To this end, PARP10 is emerging as an important player for anti-viral defence(*40*), is upregulated in response to interferon stimulation(*41*), and targets the acidic residues of Asp and Glu (*42*).

We wanted to test if Arg-C Zero could be utilized for bottom-up proteomics under mild acidic conditions which are not compatible with Trypsin digestion. To this end, we purified the catalytic domain of PARP10, and incubated this with NAD^+^ to promote ADP-ribosylation. We then used Arg-C Zero for proteolytic digestion under mild acidic conditions to preserve the labile ADPr modifications, and analysed the peptides by MS (Figure 6A). After stringent filtering to ensure confident localisation, we were able to identify 11 ADP-ribosylation sites within the ADP-ribosyl transferase (ART) domain, including several on Glu and Asp which are alkaline-labile (Figure 6). To ensure that the localisation of the ADP-ribose moiety was correct, we also annotated the diagnostic MS2 ion series of the modified peptide with the highest intensity. We found that the z4 and c10 ions uniquely demonstrate the presence of a 541.06Da mass shift corresponding to the mass of intact ADPr within the mass spectrum, confirming that Glu866 is the major auto-modification site of the ART domain of PARP10 (Figure 6C). Finally, we wanted to interrogate the modification site within the context of the structure of PARP10 ART. To achieve this, we highlighted the E866 residue of a PARP10 ART structure solved using X-ray diffraction (PDB:9FRP), and found this residue to be surface exposed, confirming its availability as an ADP-ribosylation acceptor site (Figure 6D). We also note that the residue is not in proximity to the catalytic core, which may suggest that this modification-site is established *in trans* by a partner PARP10 molecule.

**Figure 6.**
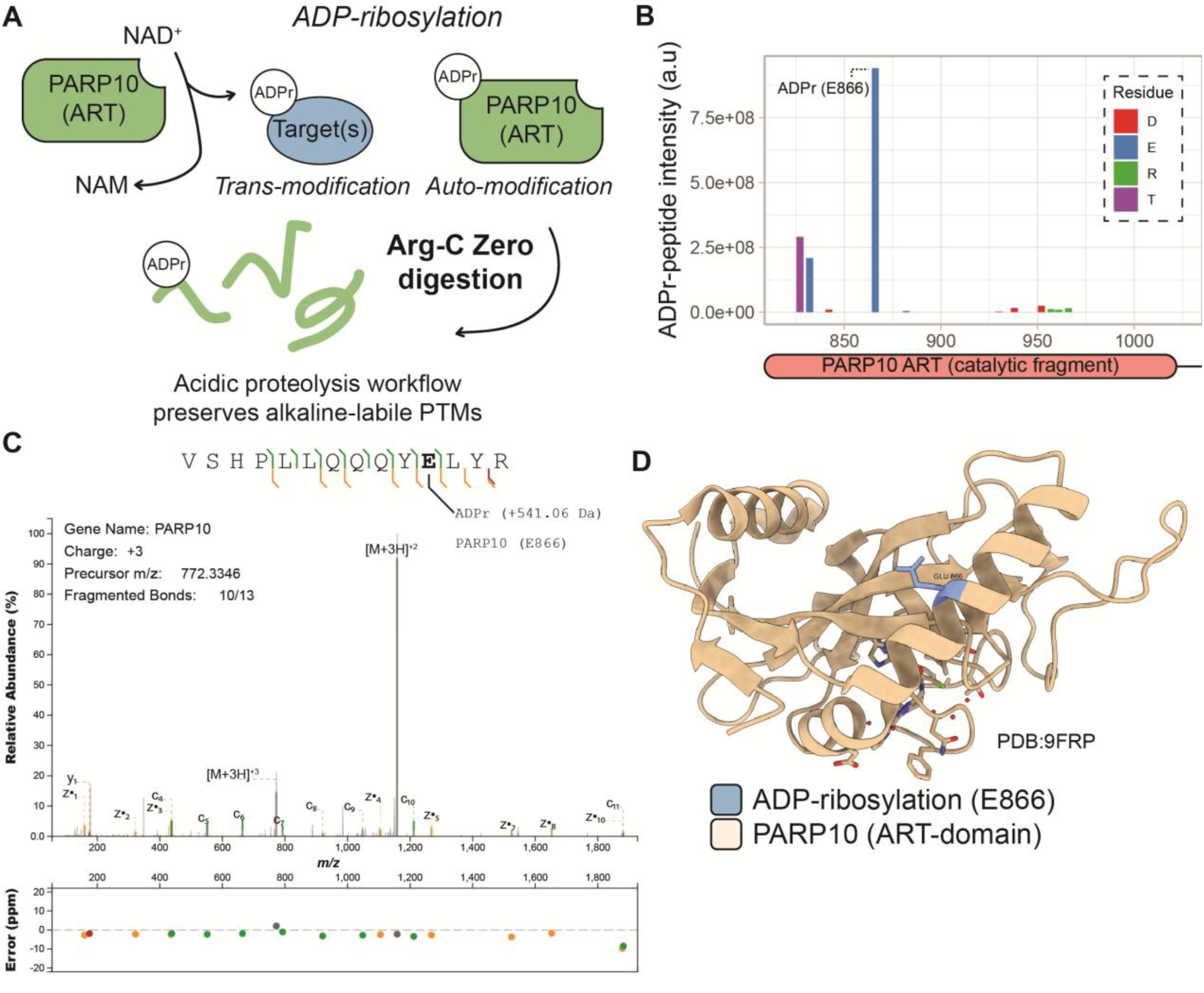
Acid compatible digestion using Arg-C Zero preserves labile PTMs. **(A)** ADP-ribosylation reaction. PARP10 utilises NAD^+^ to attach ADP-ribose (ADPr) to target proteins and itself, with nicotinamide (NAM) as a byproduct. Arg-C Zero is permissive with acidic digestion conditions which may preserve these PTMs. **(B)** Site-specific analysis of ADPr modifications on the PARP10 ADP-ribosyl transferase (ART) domain. **(C)** Annotated spectrum of the VSHPLLQQQYE(ADPr)LYR peptide, the ion series demonstrates localisation of the ADPr-modification (+541.06Da) to Glu866 within PARP10. Mass error (in ppm) of the diagnostic fragments are shown below. **(D)** Crystal structure of PARP10 ART domain (PDB:9FRP) shown in beige, with the top ADP-ribosylation site Glu866 shown in teal. A phthalazinone-based inhibitor stabilising the catalytic core is shown as a stick model.

## Discussion

The characterization of Arg-C Zero reveals several advantageous properties that position it as a valuable addition to the proteomics enzyme toolkit. The histidine-cysteine catalytic dyad, shared with the gingipain family but distinct from the serine mechanism of most commercial proteases and the cysteine mechanism of Clostripain, underpins Arg-C Zero unique biochemical profile. Structural comparison confirmed that Arg-C Zero and clostripain share no meaningful structural similarity (TM-score 0.308, Supplementary Figure 1A, and low sequence identity, Supplementary Figure 1B), consistent with convergent evolution of arginine specificity across unrelated protease families. This structural divergence implies that the distinct active site and general 3D-structure architecture of Arg-C Zero confers practical advantages, including compatibility with mild-to-strong denaturing conditions and a broad operative pH range.

Our systematic evaluation demonstrates that Arg-C Zero achieves exceptional cleavage fidelity across a wide range of digestion conditions (Figure 2). The enzyme maintained high efficiency at urea concentrations up to 4M, with optimal performance at 1 and 2M, a finding consistent with mild urea-mediated unfolding enhancing substrate accessibility under denaturing conditions (Figure 2A, B). By contrast, GuHCl was inhibitory even at 0.5M, reflecting the stronger chaotropic effect of guanidinium on the cysteine catalytic mechanism. These results provide clear practical guidance, where urea-based denaturation is fully compatible with Arg-C Zero digestion, whereas GuHCl should be removed completely prior to proteolysis. Regarding pH, the enzyme exhibited optimal performance between 6.5 and 8.5, with the lowest miss cleavages rates observed at pH 8.5, suggesting that the ionization state of the active site histidine residue is most catalytically favorable under mildly alkaline conditions (Figure 2D-F). The enzyme retained substantial activity across a twenty-fold range of enzyme-to-substrate ratios (1:25 to 1:500), with a very robust fidelity even at most diluted ratios, demonstrating practical robustness for workflows where enzyme consumption must be minimized (Figure 2G-I).

The iceLogo analysis of semi-specific cleavage events confirmed the strict arginine specificity of Arg-C Zero, with overwhelming enrichment of the P1 arginine position and no significant secondary specificities flanking positions (Figure 4A). At the population level, 98.98% of all identified peptides carried no internal arginine missed cleavages (Figure 4B). This performance substantially exceeds what is typically achieved with other enzymes, for instance, Trypsin in complex proteome digests, where miss cleavage rates of 23,4% (DDA) and 31,5% (DIA) were observed for Trypsin in our direct benchmark comparison (Figure 3G), which are consistent with commonly reported values under standard conditions (*43*). The high cleavage fidelity of Arg-C Zero simplifies database search space, reduces peptide redundancy, and improves quantification reproducibility in label-free workflows.

Our direct benchmarking against Trypsin under standard bottom-up proteomics revealed that the two enzymes are able to achieve similar and complementary proteome depth, with notable differences in terms of cleavage efficiency and peptide identifications. In DDA mode, Arg-C Zero identified fewer peptides per replicate (∼9,500 versus ∼12,400 for Trypsin), but achieved comparable protein group identifications (2,467 versus 2,333), Thus demonstrating that the arginine-specific peptide repertoire provides sufficient sequence coverage to identify a similar proteome depth despite the reduced peptide count. This is a consequence of Arg-C Zero’s longer peptide lengths compared to Trypsin (mean 15 amino acids versus 12 amino acids for Trypsin). The same trend applies to DIA mode: while Arg-C Zero generates substantially fewer peptides (∼67,000 vs ∼154,000), this reduction does not affect the protein groups identified for both enzymatic digestions (8,284 for Arg-C Zero and 8,389 for Trypsin). Venn diagrams for both acquisition modes show the great overlap in the protein detected for both conditions (Figure 3C, F).

A practical consequence of this reduced peptide complexity is the improvement in data processing time (Figure 3J). Spectronaut analysis of the Arg-C Zero raw files was 30% faster than the equivalent Trypsin datasets, explained by the smaller search space generated by Arg-C Zero cleavage specificity as well as the lower miss cleavage rates: Arg-C Zero near-complete cleavage (99,4% zero miss cleavages in DDA, and 98,9% in DIA) versus Trypsin higher miss cleavage rates (23,4% in DDA and 31,5% in DIA).

A particularly noteworthy finding is Arg-C Zero’s behaviour at proline adjacent cleavage sites. Trypsin exhibits a well-characterized efficiency deficit at R-P and K-P sequences. While proline at P1’ position was the most frequent driver of miss cleavages in our Arg-C Zero dataset, accounting for ∼22% of all miss cleavages events, the enzyme still retained 94.9% cleavage efficiency specifically at R-P bonds (Figure 4C-D). This positions Arg-C Zero as superior to Trypsin for the complete digestion of proline-rich proteins and protein regions, with practical benefits for the analysis of proline-rich proteins, where proline-flanked arginine residues are common.

For histone PTM analysis, Arg-C Zero demonstrates clear advantages over conventional tryptic digestion. While established workflows typically rely on chemical derivatization strategies to generate peptides of appropriate length for LC–MS/MS analysis, digestion with Arg-C Zero directly produces histone peptides within a suitable size range (≈ 5-15 amino acids), simplifying the workflow and reducing variability associated with derivatization efficiency. Using this enzyme, the majority of histone PTMs can be reliably detected and quantified, enabling comprehensive characterization of histone PTM patterns. Nevertheless, derivatization strategies remain necessary for certain highly hydrophilic peptides, such as those containing H3K4, which are poorly retained in standard reverse-phase LC–MS/MS workflows. Combining Arg-C Zero digestion with targeted derivatization strategies could further increase histone coverage, enabling detection of these challenging peptides while retaining the workflow simplicity.

For the analysis of alkaline-labile PTMs, there is a growing need for proteases that function with high specificity and efficiency under acidic conditions. To this end, hydrogen-deuterium exchange (HDX) MS relies on performing proteolytic digestion under strongly acidic conditions to prevent back-exchange with the solvent during proteolysis (*44*). To date, the gold standard for performing such acid-compatible digestion is with proteases like pepsin, which unfortunately has a low specificity (*45*), limiting its application for large-scale bottom-up proteomics of complex samples. Recently, the P13ase protease (*Aspergillus saitoi*) was characterized for acidic workflows (*46*), which was also shown to improve the sequence-coverage at histones. The P13ase still possesses a relatively broad sequence-specificity for its substrates, with high preference for Lys, Arg, and Leu (∼15-20%), followed by Phe, Asp, and Glu (∼5-10%). This is in contrast to Arg-C Zero, which shows unprecedented specificity towards Arg alone. While both pepsin and P13ase find important applications for acidic proteomics workflows, especially when strong acidic conditions are required for digestion (pH ∼2.5-4), Arg-C Zero could be utilized for enhanced specificity when mild acidic conditions are sufficient.

The enzyme’s broad compatibility with standard mass spectrometry buffers and sample preparation methods facilitates adoption into existing workflows without substantial protocol modifications. Arg-C Zero functions effectively in HEPES, Tris, and phosphate buffers commonly used in proteomics, however, it is worth noting that a practical consideration is the choice of reducing agent to maintain Arg-C Zero’s cysteine protease activity. While TCEP is increasingly favored over DTT in proteomics workflows due to its greater stability and odorless handling, our evaluation revealed that TCEP produces a large chromatographic artifact eluting at the tail end of chromatographic gradient, attributable to TCEP itself or its oxidized derivatives (Supplementary Figure S2B). This artifact was absent when DTT was used and was reproducible across replicate injections. Importantly, cleavage efficiency was equivalent between the two reducing agents (missed cleavages ≤0.4% for both), so the recommendation to use DTT is driven entirely by analytical compatibility rather than enzymatic performance. Users relying on TCEP in upstream steps, for example in protein extraction buffers, should therefore ensure complete buffer exchange before digestion, or substitute DTT at the digestion step. This consideration is particularly relevant for nDIA workflows analyzed on sensitive instruments such as the Orbitrap Astral, where large co-eluting peaks can suppress precursor signal and degrade quantitative performance.

Finally, proteins with few arginine residues will remain underrepresented in single-enzyme Arg-C Zero workflows; multi-enzyme strategies combining Arg-C Zero with Trypsin or Lys-C offer the most practical route to maximizing proteome depth in discovery-mode experiments.

Future developments may focus on several directions. First, combining Arg-C Zero with complementary proteases in multiplexed strategies could maximize proteome coverage while maintaining high cleavage fidelity. Second, the enzyme’s properties make it attractive for middle-down proteomics approaches, where longer peptides (20-60 residues) are analyzed to preserve information about combinatorial PTMs. The tolerance to mild denaturing conditions and resistance to modification-dependent cleavage bias could be particularly valuable for analyzing intact protein domains or investigating PTM crosstalk. Third, engineering efforts could focus on further improving stability and reducing agent requirements, perhaps through rational design of the active site or directed evolution strategies.

In conclusion, Arg-C Zero represents a high-fidelity alternative to traditional proteolytic enzymes, offering exceptional specificity for arginine residues through its histidine-cysteine catalytic mechanism. While it does not achieve the comprehensive coverage of LysC/Trypsin combinations for unbiased proteome profiling, its unique cleavage specificity enables access to protein regions that are poorly represented by tryptic approaches. The enzyme’s robust performance across varying experimental conditions, high reproducibility, and particular suitability for PTM analysis position it as a useful tool for expanding the analytical capabilities of modern proteomics laboratories. We anticipate that Arg-C Zero will find widespread application in targeted proteomics studies, PTM characterization workflows, and multi-protease strategies designed to maximize sequence coverage and analytical depth.

## Acknowledgments

We thank KPL ApS (Copenhagen, Denmark) for providing purified Arg-C Zero enzyme and technical support. Work at The Novo Nordisk Foundation Center for Protein Research (CPR) is funded in part by a donation from the Novo Nordisk Foundation (NNF14CC0001, NNF21OC0072070, and NNF24SA0098829). This project was supported by a center-of-excellence grant from the Danish National Research Foundation to Copenhagen Center for Glycocalyx Research (DNRF196). The work was also supported by PLATO, a grant from the Danish Agency of Higher Education and Science to establish the PLATO research infrastructure: Danish National Mass Spectrometry Platform for Proteomics and Biomolecular Imaging (grant no. 5229-00012B, www.sdu.dk/PLATO.

## Author Contributions

C.H.R.: Conceptualization, methodology, investigation, formal analysis, data curation, visualization, writing – original draft. T.S.B.: Conceptualization, methodology, resources, supervision, writing, review and editing. J.V.O.: Conceptualization, supervision, funding acquisition, writing, review and editing, project administration. J.D.E.: investigation, formal analysis, visualisation, and writing – review & editing. Y.L.: investigation and resources. K.P.: investigation. I.A.: supervision, funding acquisition, and writing – review & editing. E.L.B: Methodology, investigation, formal analysis, writing - review and editing. O.N.J: Supervision, funding acquisition, writing - review and editing.

## SUPPLEMENTARY FIGURES

**Supplementary Figure 1.**
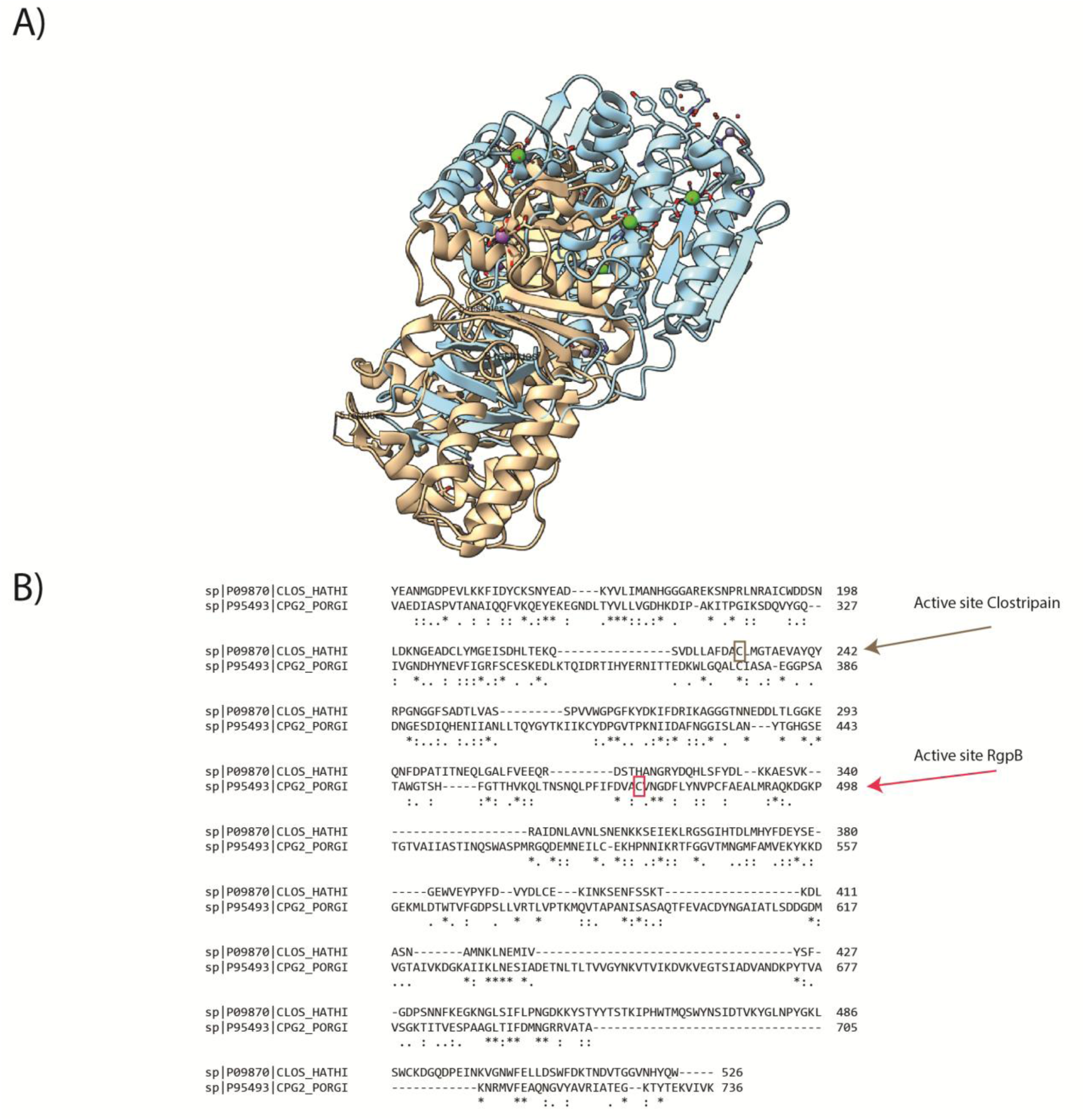
Structural and sequence comparison of Clostripain and RgpB (Arg-C Zero). **(A)** Superposition of the three-dimensional structures of Clostripain (Clostridium histolyticum, wheat; UniProt P09870) and RgpB (Porphyromonas gingivalis, blue; UniProt P95493). **(B)** Multiple sequence alignment generated with Clustal Omega (v1.2.4) of Clostripain (sp|P09870|CLOS_HATHI) and RgpB (sp|P95493|CPG2_PORGI). Arrows indicate the positions of the catalytic cysteine residues forming the His-Cys dyad in each enzyme. Conserved positions are indicated by asterisks (*), strongly similar positions by colons(:), and weakly similar positions by periods (.).

**Supplementary Figure 2.**
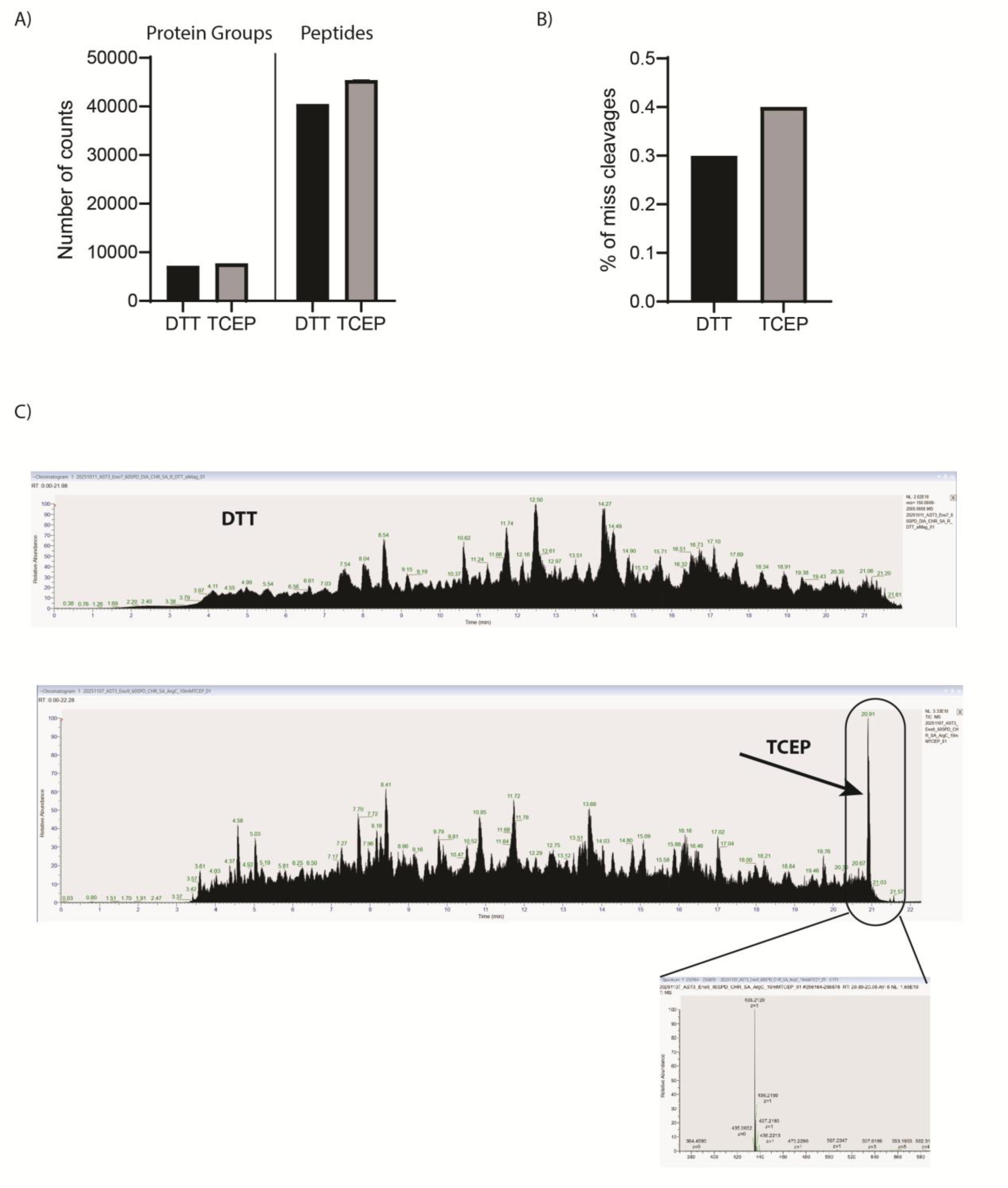
Comparison of DTT and TCEP as reducing agents for Arg-C Zero digestion and identification of a TCEP-derived contaminant ion. **(A)** Number of protein groups and peptides identified by LC-MS/MS following Arg-C Zero digestion with 10mM DTT or 10mMTCEP as reducing agent. (B) Percentage of missed cleavages observed under each condition. (C) Base peak chromatograms of DTT- (top) and TCEP-reduced (middle) samples. The TCEP chromatogram displays a prominent late-eluting peak at ∼20.9 min (RT 20.89-20.95), absent in the DTT condition. MS1 spectrum of this peak (bottom) reveals a dominant ion at m/z 435.2120 (z=1).

**Supplementary Figure 3.**
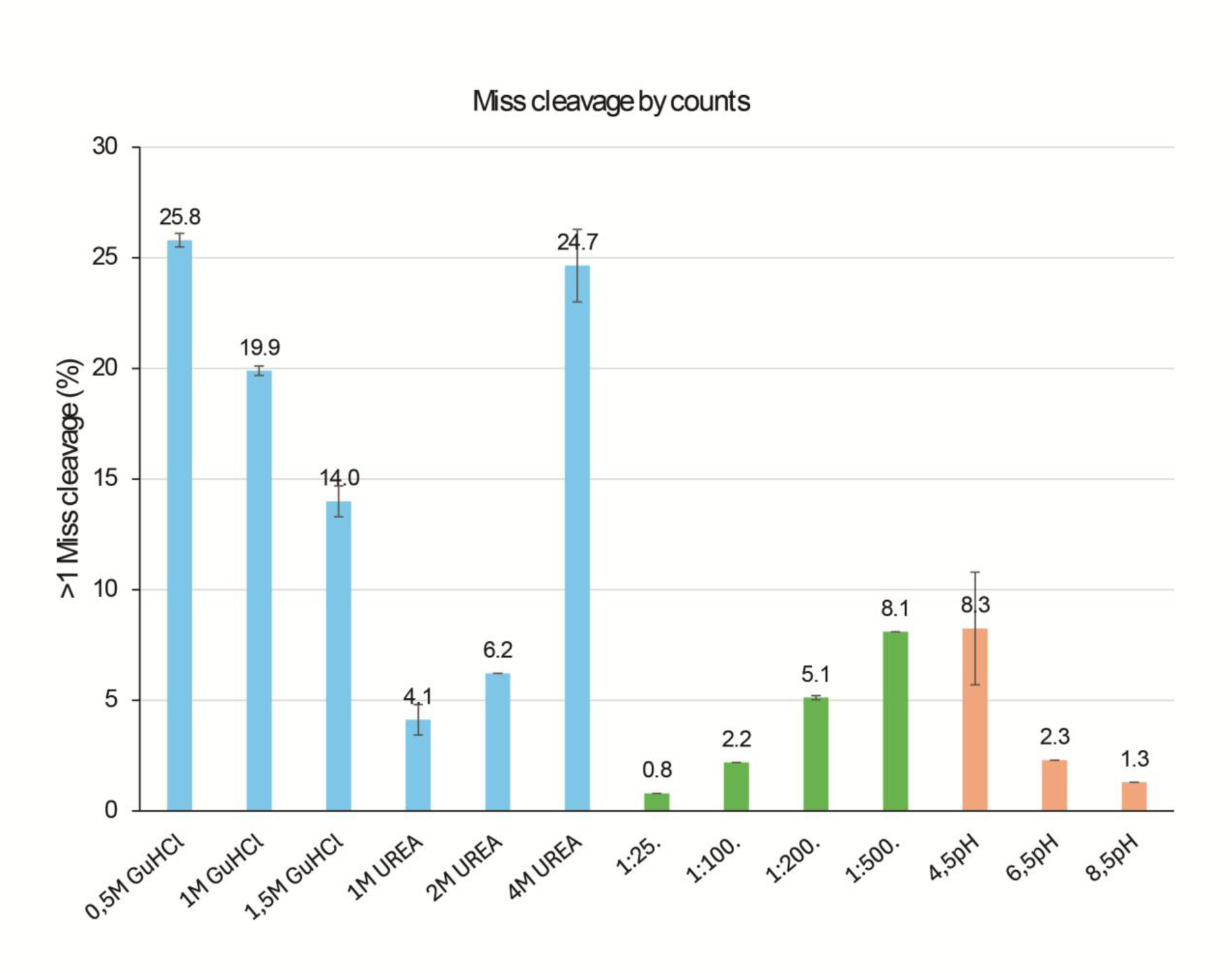
Effect of denaturant, enzyme-to-substrate ratio, and pH on Arg-C Zero digestion efficiency. Bar chart showing the percentage of peptides with more than one missed cleavage (>1 MC) across three sets of conditions: denaturant type and concentration (blue; 0.5–1.5 M GuHCI and 1–4 M urea), enzyme-to-substrate (E:S) ratio (green; 1:25 to 1:500), and digestion pH (salmon; pH 4.5, 6.5, and 8.5). Data are shown as mean± SD (n = 4).

